# Analysis of commonly expressed genes between first trimester fetal heart and placenta cell types in the context of congenital heart disease

**DOI:** 10.1101/2022.01.06.475249

**Authors:** Rebecca L. Wilson, Victor Yuan, Jennifer Courtney, Alyssa Tipler, James Cnota, Helen N. Jones

## Abstract

Congenital heart disease (CHD) is often associated with fetal growth abnormalities. During the first trimester of pregnancy, the heart and placenta develop concurrently, and share key developmental pathways. Hence, it is hypothesized that defective morphogenesis of either organ is synergistically linked. However, many studies determined to understand the mechanisms behind CHD overlook the contribution of the placenta. In this study, we aimed to identify commonly expressed genes between first trimester heart and placenta cells using two publicly available single cell sequencing databases. Using a systematic computational approach, we identified 328 commonly expressed genes between heart and placenta endothelial cells and enrichment in pathways including Vasculature Development (GO:0001944, FDR 2.90E-30), and Angiogenesis (GO:0001525, FDR 1.18E-27). We also found, in comparison with fetal heart endothelial cells, 197 commonly expressed genes with placenta extravillous trophoblasts, 128 with cytotrophoblasts and 80 with syncytiotrophoblasts, and included genes such as *FLT1, GATA2, ENG* and *CDH5*. Finally, comparison of first trimester cardiomyocytes and placenta cytotrophoblasts revealed 53 commonly expressed genes and enrichment in biological processes integral to cellular function including Cellular Respiration (GO:0045333; FDR 5.05E-08), Ion Transport (GO:0006811; FDR 2.08E-02), and Oxidation-Reduction Process (GO:0055114; FDR 1.58E-07). Overall, our results identify specific genes and cellular pathways common between first trimester fetal heart and placenta cells which if disrupted may concurrently contribute to the developmental perturbations resulting in CHD.

## Introduction

Congenital heart disease (CHD) is the most common birth defect ^1^. Affecting approximately 1% of live births, it is the leading cause of infant mortality related to birth defects ^2^, and is often the result of perturbations in normal programming of cardiac development. In approximately one third of babies, the CHD is classified as severe and requires intervention in the first year of life ^3^. Survival of such interventions is hindered by the fact that there is a higher incidence of fetal growth restriction (FGR) and preterm birth in CHD pregnancies. Such associations suggest that the perturbations to cardiac programming are also affecting placental development, however, the etiology of fetal growth abnormalities in CHD is largely unknown.

*In utero* the placenta and heart develop concurrently ^4-8^. Placenta and heart development also share key developmental pathways; hence it is reasonable to assume that deleterious changes in gene expression during first trimester development of one are likely to perturb morphogenesis of the other ^9^. The heart is the first organ to develop within the fetus with cells destined to become the mature heart originating from the mesoderm ^10^. The asymmetrical nature of heart development at both the organ and tissue level contribute to the complex nature of the molecular events which occur in the first trimester. At the same time as the early heart is developing, vascular development of the placenta is also occurring ^11^. Arising from the trophectoderm and extra-embryonic mesoderm, trophoblastic structures branch and give rise to the primary and secondary placental villi. Differentiation of the mesenchymal cells inside the villi eventually result in the first hemangiogenic precursor cells and ultimately develop into the placental vasculature.

In both heart and placental development, the cellular and signaling events that occur during the first trimester will ultimately determine the fate of the developing heart and placental vasculature. Numerous animal models have shown that disrupted expression of genes associated with common vasculogenesis and angiogenesis pathways result in both placental and cardiac defects ^12^. In addition to molecular signals which modulate heart development in the first trimester, mechanical forces are also required. For example, there is evidence which suggests that ventricular wall expansion in the heart switches from embryonic to placental control after the onset of maternal blood flow into the placenta ^13^. Thus, there is the potential in pregnancies complicated by diseases such as preeclampsia or FGR, where there is inadequate or changed placental blood flow, that a synergistic effect with placental insufficiency exacerbates the developmental defect in the heart ^9^.

The parallel development of the placenta and heart *in utero* has led to the hypothesis that defective morphogenesis of either organ is synergistically linked. Indeed, there are several mutant mouse models which exhibit both defective heart and placenta development ^9^. In some cases, the heart defects were shown to be secondary to defective placental development, possibly as a consequence of insufficient placental blood flow affecting early embryonic cardiac function ^14^. However, despite the reoccurring link between heart and placental defects, studies focused on CHDs rarely acknowledge the potential contribution of the placenta. It is estimated that over 300 genes collectively contribute to CHDs in humans ^15^, and studies using conditional knockout animal models have confirmed that disrupting many of these genes leads to cardiac defects ^16^. In these studies, conditional knockout occurred in cardiac tissue only, however, many of the genes are known to also be expressed in the placenta. Therefore, there is a heightened need to identify and better understand common gene expression profiles between the heart and placenta, particularly in the first trimester, in order to improve the understanding of the mechanisms by which CHDs occur.

Advances in single-cell sequencing techniques are now allowing for greater in-depth analyses of how a given gene is expressed in different organs like the placenta and heart. In this study, we utilize two publicly available databases analyzing gene expression in first trimester human placenta ^17^ and fetal heart cells ^18^. Using a systematic, computational approach, we aimed to identify commonly expressed genes between first trimester placenta endothelial cells (ECs), extravillous trophoblasts (EVTs), cytotrophoblasts (CTBs) and syncytiotrophoblasts (STBs), and first trimester fetal heart ECs and cardiomyocytes. Our results identify specific genes and cellular pathways common between placental and fetal heart cells which if disrupted may concurrently contribute to the developmental perturbations resulting in CHD.

## Results and Discussion

Using publicly available databases ^17,18^, we compared single-cell gene expression profiles from first trimester human embryonic heart ECs and cardiomyocytes with first trimester human placental ECs, EVTs, CTBs and STBs. Principal component analysis based on global gene expression showed clustering of fetal heart and placenta ECs, whilst EVTs, CTBs, and STBs also clustered together (Figure 1). Compared to heart ECs, there was 328 commonly expressed genes with placental EC, 197 with EVTs, 128 with CTBs and 80 with STBs (Figure 1). For heart cardiomyocytes, there was 16 commonly expressed genes with placental endothelial cells, 36 genes in common with EVTs, 53 genes with CTBs and 22 commonly expressed genes with STBs (Figure 1). Full details of commonly expressed genes and pathway enrichment analysis can be found in Supplemental Material.

**Figure 1.**
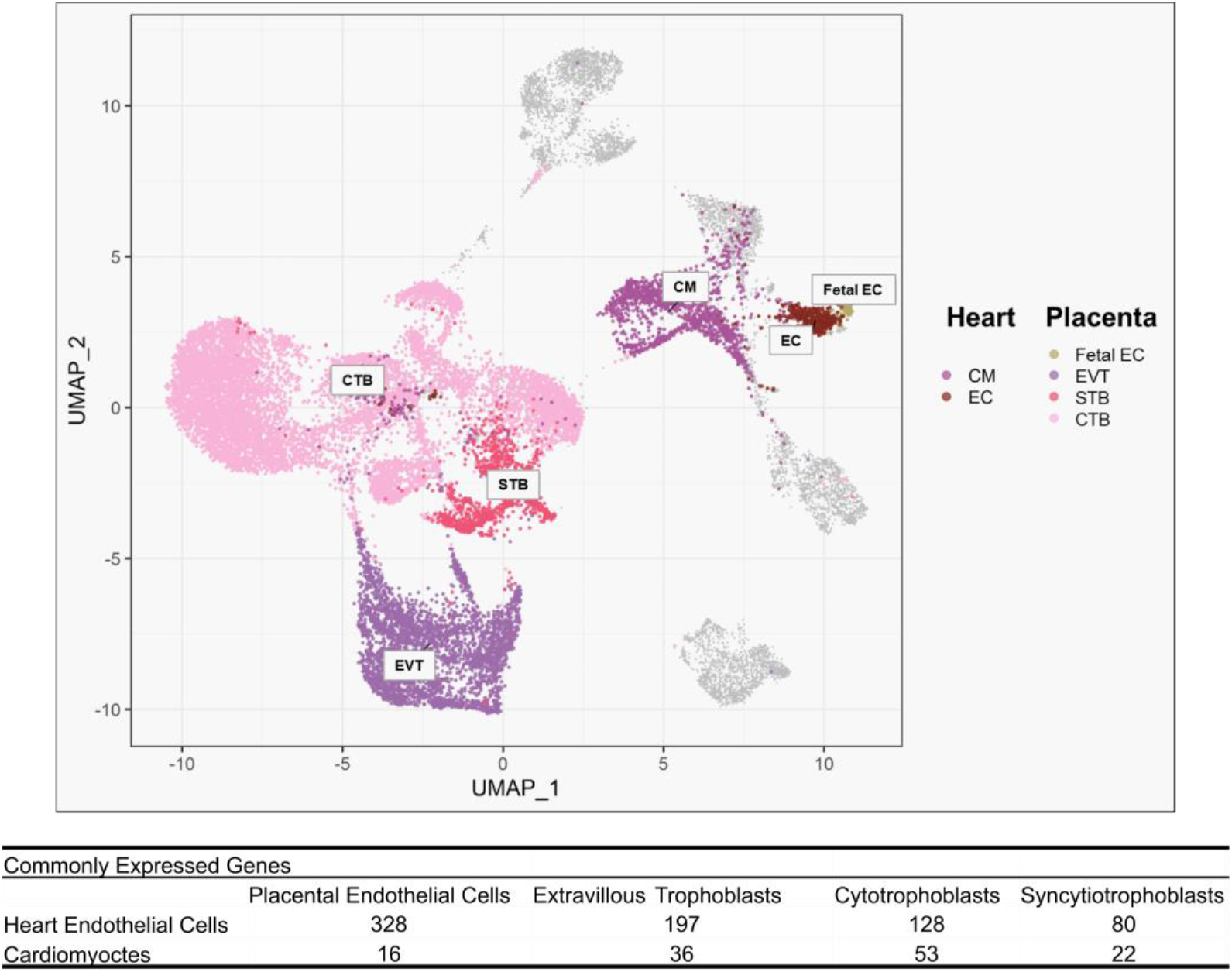
Principal component analysis of global gene expression in first trimester heart and placenta cell types and number of commonly expressed genes between heart endothelial cells (EC) or cardiomyocytes (CM) and placenta (fetal) EC, extravillous trophoblasts (EVT), syncytiotrophoblasts (STB) and cytotrophoblasts (CTB).

### Commonly expressed genes between first trimester placental and fetal heart endothelial cells

As expected, heart and placental endothelial cells had the most commonly expressed genes. Furthermore, functional enrichment analysis using ToppGene – ToppFun revealed enrichment of genes associated with biological functions including *Vasculature Development* (GO:0001944, FDR 2.90E-30), *Angiogenesis* (GO:0001525, FDR 1.18E-27), *Heart Development* (GO:0007507, FDR 9.15E-12) and *In utero Embryonic Development* (GO:0001701 FDR 6.84E-07)(Figure 2a), reflecting the role of endothelial cells in both the placenta and heart. Early in heart development, the primitive heart consists of 2 cardiac progenitor cell layers: endocardial endothelial cells and cardiomyocytes ^19^. Originating from the rostrolateral mesoderm, it is the endocardial endothelial cells which differentiate and give rise to the other cell types of the heart including cells in the cardiac valves and chambers ^20^. Similarly in the placenta, precursor endothelial cells are derived from the mesoderm ^21^. Vasculogenesis begins within the first 18-20 days after conception ^22^. Precursor endothelial cells form vessels beneath the trophoblastic epithelium through a combination of cell replication and stromal cell recruitment, with villous circulation formed by 6 weeks post-conception. As pregnancy progresses, there is expansion of the fetal capillary bed via both branching and non-branching angiogenesis in order to fully support rapid growth of the fetus in late pregnancy ^7^.

**Figure 2.**
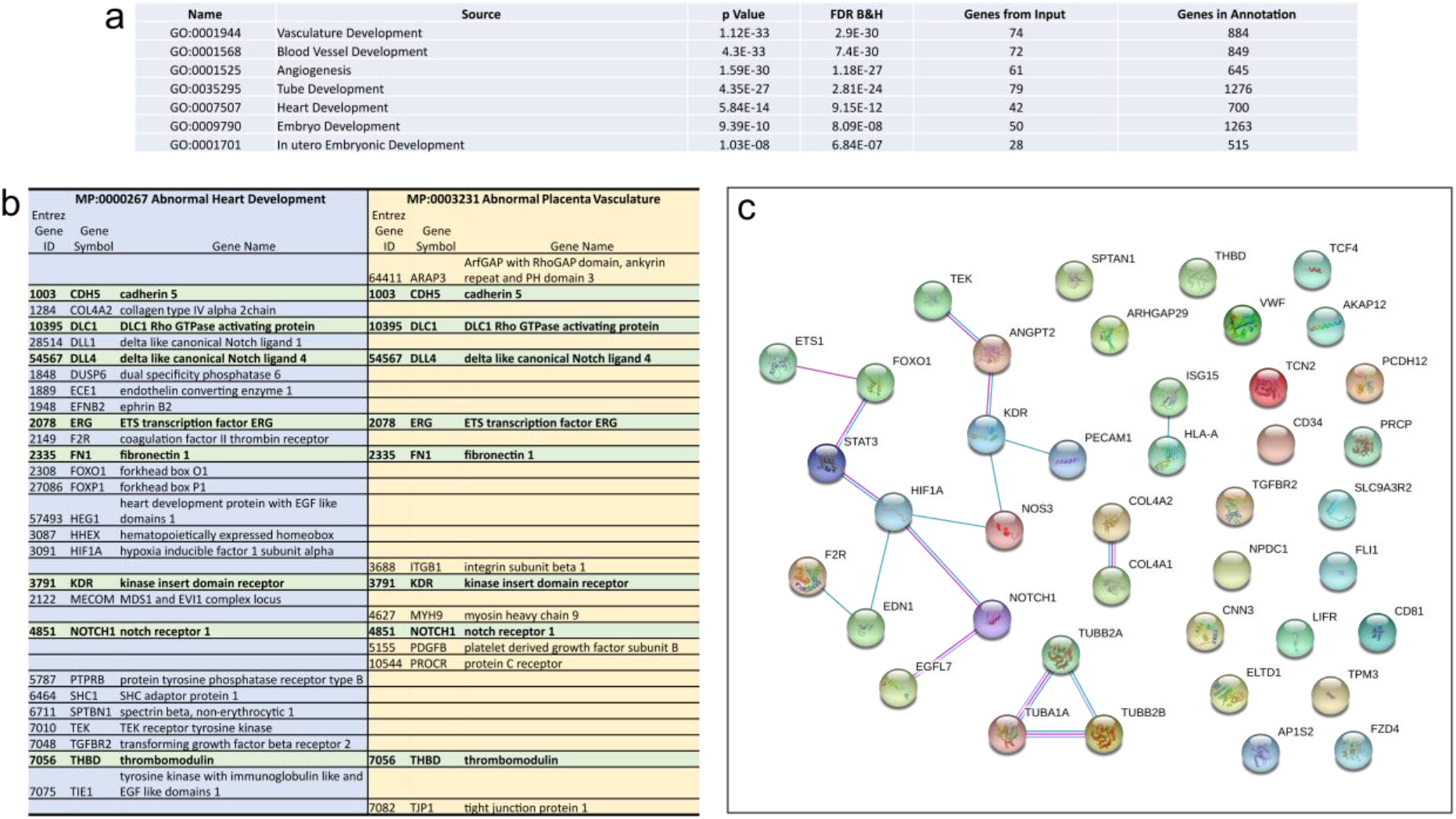
Comparison of commonly expressed genes between first trimester placenta and heart endothelial cells. **a**. Enrichment analysis of GoBiological processes of the commonly expressed genes. **b**. Comparison of commonly expressed genes between placenta and heart endothelial cells associated with abnormal heart development and placenta vasculature in mouse phenotypes. **c**. Protein-protein interactions of 42 commonly expressed genes between placenta and heart endothelial cells associated with human congenital diseases.

We also assessed commonly expressed genes between heart endothelial cells and placental endothelial cells for enrichment in mouse phenotypes. This revealed enrichment for phenotypes such as *Abnormal Heart Development* (MP:0000267 FDR 7.27E-05) and *Abnormal Placental Vasculature* (MP:0003231 FDR 4.92E-04). Moreover, there was enrichment in numerous mouse phenotypes associated with abnormal placental development and function including *Embryonic Growth Retardation* (MP:0003984 FDR 6.25E-06), *Lethality throughout Fetal Growth and Development* (MP:0006208 FDR 1.25E-08) and *Abnormal Visceral Yolk Sac Morphology* (MP:0001718 FDR 2.96E-10). Comparison of the 26 genes associated with abnormal heart development and abnormal placental vasculature showed 8 genes similar to both phenotypes including *notch receptor 1* (*NOTCH1*) and its receptor ligand *DLL4* (Figure 2b). Notch signaling is highly conserved and provides a means by which cells can influence neighboring cells through receptor-ligand binding ^23^. In the heart, Notch signaling has been shown to regulate cardiac cell fate and orchestrate cardiac chamber and valve morphogenesis ^24^. Additionally, there are several Notch pathway genes in which mutations have been implicated in CHDs including *NOTCH1, NOTCH2, DLL4, JAG1* and *MAML2* ^15^. Mutations in *NOTCH1* are associated with numerous CHDs ranging from issues with development of the bicuspid aortic valve to HLHS ^25^.

In terms of the placenta, Notch1^-/-^ knockout mice are embryonically lethal at gestational day 11.5 ^26^. Analysis of the placenta has shown that whilst fusion of the allantois with the chorionic plate occurs, fetal blood vessels do not form within the labyrinthine region of the placenta ^27^. This phenotype has also been reported in a mouse model where Notch1 was conditionally knockout only in the endothelium ^28^ and as such, indicates a major role for Notch1 in placental development.

From a translational perspective, there was enrichment in genes commonly expressed between placental and heart ECs for *Human Congenital Abnormalities* (C0000768 FDR 5.43E-06). The 42 genes included a specific cluster of genes, including *NOTCH1, HIF1A, KDR*, and *NOS3*, in which there are known protein-protein interactions (Figure 2c). *NOS3*, or endothelial nitric oxide synthase is one of three nitric oxide synthase isoforms and is crucial in the regulation of vascular integrity and homeostasis. In pregnancy, the importance of *NOS3*, particularly towards placenta angiogenesis and vascular development is well established ^29^. *Nos3* knockout mice are characterized with fetal growth restriction during pregnancy and placental dysfunction ^30,31^. Additionally, genetic variants in *NOS3* gene have been associated with increased risk of the pregnancy disease preeclampsia ^32^. In terms of cardiac development, *NOS3* is known to play an important role ^33,34^ and polymorphisms in *NOS3* have been shown to be associated with increased risk of sporadic CHD, and more specifically a 62% increased risk of perimembranous ventricular septal defects ^35^.

### Commonly expressed genes between first trimester heart endothelial cells and first trimester placenta trophoblasts

Early placental development and function is determined by the various groups of trophoblasts; disrupted cellular signaling events within these trophoblast cells can lead to defective placentation and the development of obstetric diseases. There were 197 commonly expressed genes between fetal heart ECs and placental EVTs and included enrichment in biological functions such as, *Vascular Development* (GO:0001944, FDR 2.03E-05), *Angiogenesis* (GO:0001525, FDR 1.83E-05) as well as, *Cell Migration* (GO:0016470 FDR1.21E-05), *Cell Adhesion* (GO:0007155 FDR 2.26E-05) and *Extracellular Structure Organization* (GO:0043062 FDR 1.25E-07)(Figure 3a). Enrichment in such pathways is unsurprising given the roles of both heart ECs and EVTs in their respective organs. Coinciding with the time in pregnancy in which the early heart patterning is occurring, the placental EVTs invade the maternal uterine decidua, transforming the uterine spiral arteries in preparation for the onset of placental blood flow ^36^. Numerous obstetric diseases including FGR and preeclampsia are characterized by having abhorrent maternal spiral artery transformation, most likely due to inappropriate functioning of the EVTs ^37^. Large human cohort studies have also confirmed strong positive associations between congenital heart defects and obstetric diseases like preeclampsia, preterm birth and FGR ^38-42^ further strengthening the hypothesis that the underlying pathophysiology of CHDs and placental-related pregnancy complications originate from similar biological insults.

**Figure 3.**
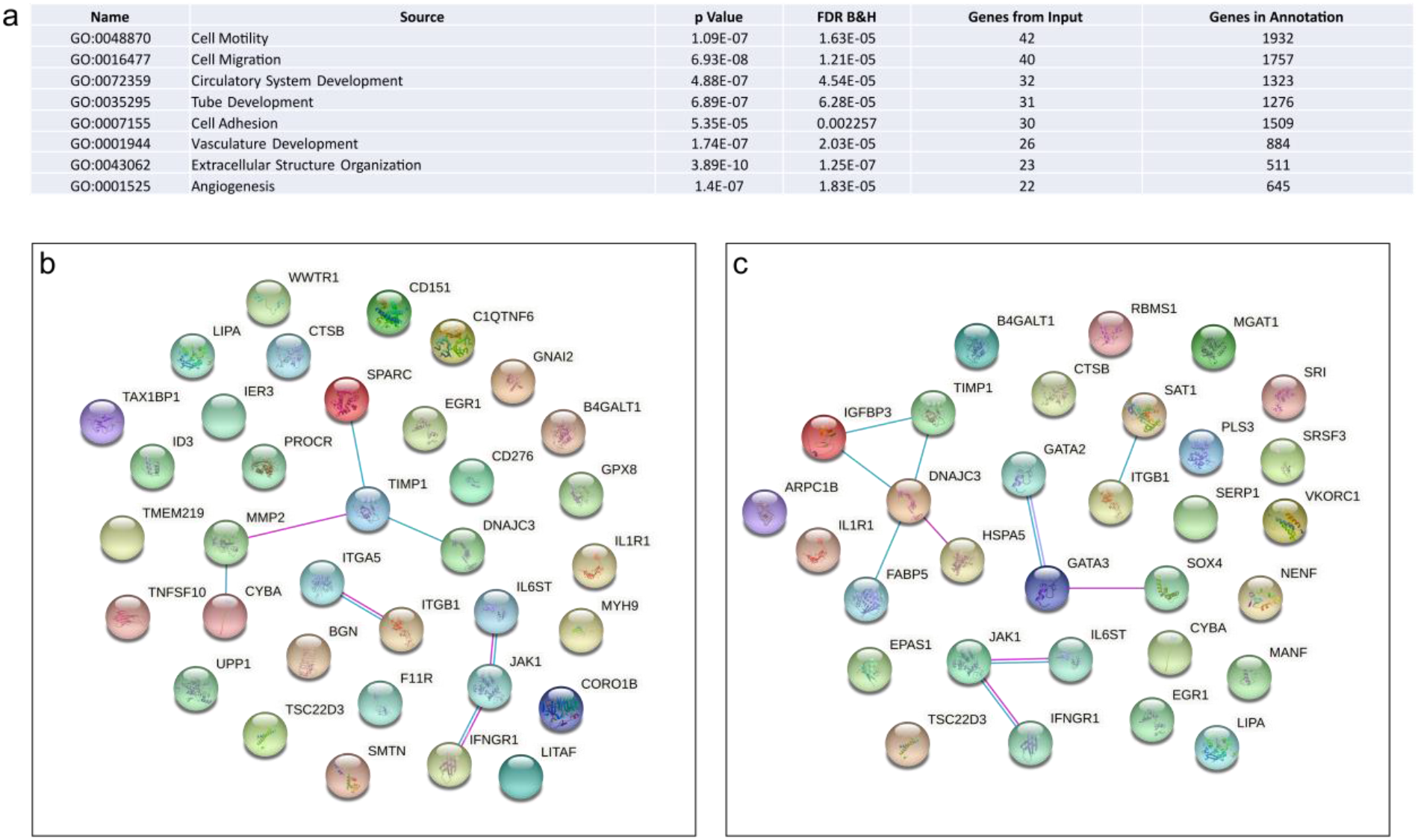
Comparison of commonly expressed genes between fetal heart endothelial cells and placenta extravillous trophoblast cells. **a**. GoBiological processes enrichment analysis of the commonly expressed genes. **b**. Protein-protein interactions of commonly expressed genes associated with abnormal hormone levels. **c**. Protein-protein interactions of commonly expressed genes associated with abnormal inflammation response.

In terms of mouse phenotypes, there was enrichment in the commonly expressed genes between fetal heart ECs and EVTs for *Abnormal Vascular Development* (M:0000259 FDR 3.27E-05) and *Abnormal Cardiovascular System Physiology* (M:0001544 FDR 9.66E-05) and included genes such as *MMP2, FN1, ENG* and *EPAS1*. Interestingly, there was also enrichment for genes associated with *Abnormal Hormone Levels* (MP:0003953 FDR 1.55E-02) and included *SOX4, GATA2*, and *GATA3* which are known to interact at a protein level (Figure 3b), as well as *Abnormal Inflammatory Response* (MP:0001845 FDR1.55E-05) including genes *JAK1, IL1R1*, and *IFNGR1* (Figure 3c). The role of the immune system in placental development and function is well established with the invading EVTs communicating with the resident uterine immune cells to invade and transform the uterus appropriately ^43^. Improper control of EVT invasion is a hallmark characteristic of placenta accreta disorder, in which EVT invasion extends beyond the maternal decidua and in severe cases, into other organs like the bowel or bladder and more is being understood about the role of immune cells in allowing such over invasion to occur ^44^. Both over- and under-invasion of the EVTs results in significant changes to placental hemodynamics and the flow of maternal blood into the placenta. It is known that early cardiac development is dependent on both genetic and environmental factors, and hemodynamic forces associated with blood flow play an important role ^45,46^. Experimentally induced alterations in hemodynamics of the umbilical vein and umbilical artery have been shown to trigger detrimental growth and remodeling cascades eventuating in major cardiac defects ^47^. Such findings support the idea that impaired maternal blood flow to the placenta, as well as genetic factors, can have a significant effect on early embryonic cardiac development and help explain why there is a strong association between placental-related pregnancy complications and CHDs.

Whilst the functional similarities between fetal heart ECs and placenta CTBs and STBs are not as well defined as between fetal heart ECs and placenta ECs and EVTs, there was still a large number of commonly expressed genes. Enrichment analysis of the 128 commonly expressed genes between fetal heart ECs and placental CTBs revealed biological processes involved in cellular function, such as, *Exocytosis* (GO:0006887 FDR 1.71E-02), *Glycoprotein Metabolic Process* (GO:0009100 FDR 8.07E-03), *Membrane Fusion* (GO:0061025 FDR 2.33E-02), *Response to Endogenous Stimuli* (GO:0009719 FDR 1.14E-02) and *Vesicle Organization* (GO:0016050 FDR 2.33E-02) as opposed to vasculature development (Figure 4a). On the other hand, commonly expressed genes between fetal heart ECs and STBs were enriched for biological processes including *Angiogenesis* (GO:0001525 FDR 3.76E-05), *Heart Morphogenesis* (GO:0003007 FDR 1.31E-02) *and Tube Morphogenesis* (GO:0035239 FDR 2.46E-05), as well as functions like *Secretion* (GO:0046903 FDR 5.90E-04) and *Organelle Fusion* (GO:0048284 FDR 2.00E-04) more typically associated with STB function (Figure 4b). There was also enrichment in mouse phenotypes associated with *Embryonic Lethality During Organogenesis, complete penetrance* (MP:0011098 FDR 5.50E-03) and *Abnormal Heart Morphology* (MP:0000266 FDR 1.14E-02) and included genes such as *FLT1, GATA2, ENG* and *CDH5* which were common to both phenotypes (Figure 4c). Analysis of protein-protein interactions between the 80 commonly expressed genes revealed a unique cluster of 6 genes hypothesized to be involved in the innate immune response (Figure 4d).

**Figure 4.**
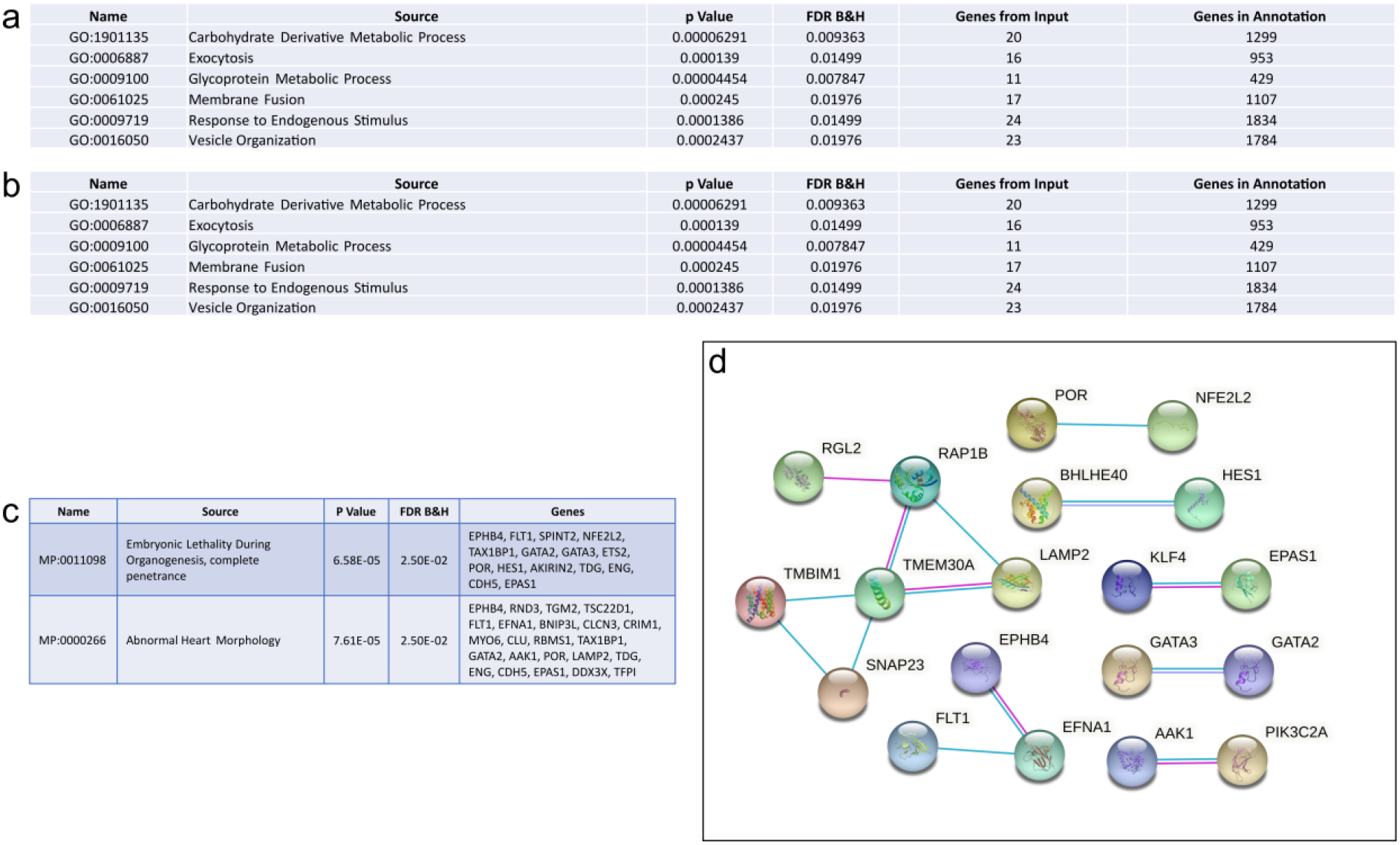
Comparison of commonly expressed genes between fetal heart endothelial cells and placenta cytotrophoblasts and syncytiotrophoblasts. **a**. GoBiological processes enrichment analysis of commonly expressed genes between heart endothelial cells and placenta cytotrophoblasts. **b**. GoBiological processes enrichment analysis of commonly expressed genes between heart endothelial cells and placenta syncytiotrophoblasts. **c**. Enrichment analysis of commonly expressed genes between heart endothelial cells and syncytiotrophoblasts in mouse phenotypes. **d**. Protein-protein interaction of the 80 commonly expressed genes between first trimester fetal heart endothelial cells and placenta syncytiotrophoblasts.

### Commonly expressed genes between first trimester cardiomyocytes and first trimester placenta trophoblasts

Despite overall gene expression profiles clustering first trimester fetal cardiomyocytes close to first trimester placenta ECs, there were only 16 commonly expressed genes between the two cell types and included the glucose transporter *SLC2A1, AK1, CAV1*, and *BEX1*. Similarly, there were very few commonly expressed genes between fetal heart cardiomyocytes and placenta EVTs and STBs: see supplemental material for detailed list of commonly expressed genes. However, comparison of fetal heart cardiomyocytes and placenta CTBs; both cell types are progenitor cells in their respective organs, revealed 53 commonly expressed genes including *BMP7, SLC38A1, GJA1*, and *DSP* (Figure 5a). These 53 genes were enriched for biological processes integral to cellular function including *Cellular Respiration* (GO:0045333; FDR 5.05E-08), *Ion Transport* (GO:0006811; FDR 2.08E-02), and *Oxidation-Reduction Process* (GO:0055114; FDR 1.58E-07)(Figure 5b). In the heart, cardiomyocytes are responsible for driving heart contraction, maturing from fetal cardiomyocytes to adult cardiomyocytes in order to sustain cycles of contraction and relaxation ^48^. Cardiomyocyte regeneration occurs naturally through proliferation of existing cardiomyocytes ^49^, although this proliferative capacity only exists during fetal development and is quickly lost after birth ^50^. In the placenta, CTBs undergo asymmetrical division where by one daughter cell re-populates the progenitor pool whilst the other differentiates and fuses with the overlying STB layer ^51^. For both cardiomyocytes and CTBs, maladaptive responses in differentiation and function are characteristic of pathological conditions including hypertension, myocardial infarction, and cardiac fibrosis in the heart ^52^, and preeclampsia and FGR in the placenta ^53^.

**Figure 5.**
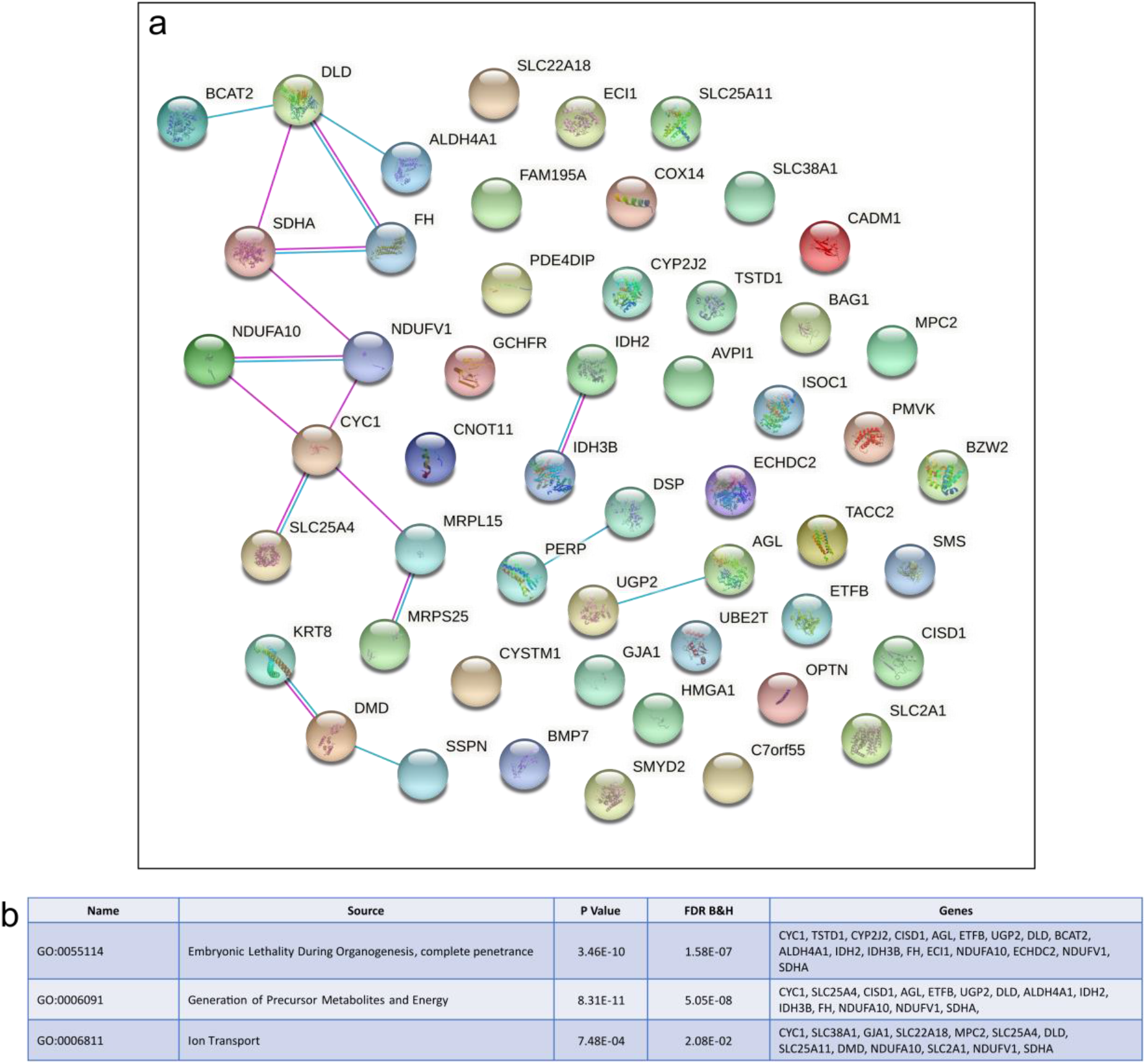
Comparison of commonly expressed genes between first trimester fetal heart cardiomyocytes and placenta cytotrophoblasts. **a**. Protein-protein interactions of the commonly expressed genes between heart cardiomyocytes and placenta cytotrophoblasts. **b**. GoBiological processes enrichment analysis of commonly expressed genes between heart cardiomyocytes and placenta cytotrophoblasts.

## Conclusion

To date, efforts to understand the mechanistic origins of congenital heart defects have largely ignored the impact of the placenta. Given the heart and the placenta develop concurrently in early gestation, and the direct physiological connection, there is the potential for investigations that ignore the role the placenta in the development of CHDs to miss crucial causative steps. Systematic analysis of single-cell gene expression profiles between first trimester heart and placenta cell types has revealed commonly expressed genes and biological pathways that are essential for normal cell and organ function and known to be associated with both CHDs and placenta-related obstetric diseases. Hence, providing further evidence that future research into the mechanisms behind CHD development need to acknowledge the contribution of the placenta.

## Methods

### Data Preparation

Single cell sequencing data was obtained from two publicly available repositories: Cui *et al*. GEO GSE106118 and Vento-Tormo *et al*. ebi ArrayExpress (E-MTAB-6701 and E-MTAB-6678). The Vento-Tormo data was downloaded as filtered, unnormalized counts. The Cui data was downloaded as already TPM-normalized. To account for the different scales of these data, both datasets were separately normalized with a natural log transformation, and then batch correction (canonical correlation analysis) was applied using the R package Seurat ^54^.

### Shared genes between first trimester heart and placenta cells

As a first step, the R package Seurat was used to identify cell-specific differentially expressed genes in a tissue-specific manner. Using a Wilcoxon Rank Sum test and a one-versus-all design, the mean expression of a gene for one cell type was compared to the mean expression of all other cell types. To define cell-specific differential expression, stringent statistical thresholds were used to minimize the proportion of false positives: the minimum log-fold change was set to >0.2, Bonferroni-adjusted p-value of <0.01, and the difference in the percentage of cells showing expression of a gene between the cell type of interest and all other cell types was >5%. To find commonly expressed genes, the cell-specific differentially expressed genes were overlapped in a pairwise fashion between heart and placenta cell types.

### Analysis of shared genes between first trimester heart and placenta cells

The commonly expressed genes between cell types of interest were entered into ToppFun (ToppGene Suite V25: ^55^) for enrichment analysis of GO Biological Process, Mice Phenotypes and Human Diseases. P values were calculated using the Hypergeometric Probability Mass Function and false discovery rate corrected using Benjamini-Hochberg methods. Potential protein interactions of shared genes of interest between cell types were analyzed using STRING (Database V11.0 ^56^). Only known protein-protein associations from curated databases and/or experimentally determined were assessed.

## Acknowledgments

This study was funded by National Institutes Health award R01HD091527 (HNJ).

## Author contributions

RLW designed the analyses, interpreted the data and wrote and drafted the manuscript. VY analyzed and interpreted the data and edited the manuscript. JC designed the analyses and edited the manuscript. AT interpreted the data and edited the manuscript. JC conceived the work and edited the manuscript. HNJ conceived the work, designed the analyses, obtained the funding and edited the manuscript. All authors have reviewed and approved the submitted version.

## Competing Interests

the authors declare no conflicts of interest

## Data Availability

Single cell sequencing data can be obtained from two publicly available repositories: GEO (GSE106118) and ebi ArrayExpress (E-MTAB-6701 and E-MTAB-6678).

## Notes

### Competing Interest Statement

The authors have declared no competing interest.

